# Population genomic and historical analysis reveals a global invasion by bridgehead processes in *Mimulus guttatus*

**DOI:** 10.1101/2020.06.26.173286

**Authors:** Mario Vallejo-Marín, Jannice Friedman, Alex D. Twyford, Olivier Lepais, Stefanie M. Ickert-Bond, Matthew A. Streisfeld, Levi Yant, Mark van Kleunen, Michael C. Rotter, Joshua R. Puzey

**Affiliations:** Biological and Environmental Sciences, University of Stirling. Stirling, Scotland, United Kingdom. FK9 4LA; Biology Department, Queen’s University. Kingston, Ontario, Canada. K7L 3N6; School of Biological Sciences, University of Edinburgh. Edinburgh, Scotland, United Kingdom. EH9 3FL; Royal Botanic Garden Edinburgh, 20a Inverleith Row, Edinburgh EH3 5LR; INRAE, Univ. Bordeaux, BIOGECO, Cestas, France. F-33612; Herbarium (ALA), University of Alaska Museum of the North, University of Alaska Fairbanks, Fairbanks, Alaska. United States of America. 99775; Institute of Ecology and Evolution, University of Oregon. Eugene, Oregon, United States of America. 97403; Future Food Beacon and School of Life Sciences, University of Nottingham, Nottingham NG7 2RD, United Kingdom; Department of Biology, University of Konstanz. Konstanz, Germany. D-78457; Zhejiang Provincial Key Laboratory of Plant Evolutionary Ecology and Conservation, Taizhou University, Taizhou, China. 318000; Department of Biological Sciences, Northern Arizona University. Flagstaff, Arizona, United States of America. 86011; Biology Department, College of William and Mary. Williamsburg, Virginia, United States of America. 23185

**Keywords:** Admixture, Approximate Bayesian Computation, bridgehead invasion, *Erythranthe*, genotype-by-sequencing, hybridisation, multiple origins, naturalisation

## Abstract

Humans are transforming species ranges worldwide. While artificial translocations trigger biological invasions with negative effects on biodiversity, invasions provide exceptional opportunities to generate ecological and evolutionary hypotheses. Unfortunately, imperfect historical records and exceedingly complex demographic histories present challenges for the reconstruction of invasion histories. Here we combine historical records, extensive worldwide and genome-wide sampling, and demographic analyses to investigate the global invasion of yellow monkeyflowers (*Mimulus guttatus*) from North America to Europe and the Southwest Pacific. By sampling 521 plants from 158 native and introduced populations genotyped at >44,000 loci, we determined that invasive North American *M. guttatus* was first likely introduced to the British Isles from the Aleutian Islands (Alaska), followed by rapid admixture from multiple parts of the native range. Populations in the British Isles then appear to have served as a bridgehead for vanguard invasions worldwide into the rest of Europe, New Zealand and eastern North America. Our results emphasise the highly admixed nature of introduced *M. guttatus* and demonstrate the potential of introduced populations to serve as sources of secondary admixture, producing novel hybrids. Unravelling the history of biological invasions provides a starting point to understand how invasive populations adapt to novel environments.

## Introduction

Increasing global connectivity is leading to widespread species translocations (Chapman, Purse, Roy, & Bullock, 2017). Most biological communities now include introduced members that have recently moved beyond their native ranges, often with negative impacts (Pysek et al., 2012; Seebens et al., 2017; Seebens et al., 2015; van Kleunen, Dawson, et al., 2015; Vila et al., 2011). Finding the origins of invaders helps develop strategies for prevention, management and eradication (Hufbauer, 2004; Hulme et al., 2008). It is also crucial for understanding to what extent invaders adapted to novel environments, along with the mechanisms of such adaptations (Dlugosch & Parker, 2008; Welles & Dlugosch, 2019).

Tracing the migration and spread of invasives is typically very challenging. Inferring introduction histories is often accomplished using historical records, genetic analyses, or a combination of both (Estoup & Guillemaud, 2010; Lombaert et al., 2010; van Boheemen, Atwater, & Hodgins, 2019). In most cases, historical records of first introduction are unavailable or unreliable. Genetic data has greatly improved our ability to study the origins of invasions, and often uses information derived from extant populations (Welles & Dlugosch, 2019). However, genetic inferences are usually confounded by demographic processes that shape the introduced populations, including multiple introduction events, bottlenecks, evolution in the introduced range, admixture and hybridisation (Bock et al., 2015; Dlugosch, Anderson, Braasch, Cang, & Gillette, 2015; Estoup & Guillemaud, 2010).

Here we use historical and genomic data to generate and test hypotheses in order to unravel the rapid worldwide invasion by the common yellow monkeyflower, *Mimulus guttatus* Fischer ex DC. (*Erythranthe spp*. (L.) G. L. Nesom; Phrymaceae), a herbaceous plant native to Western North America that was introduced across the world in the 19_th_ century (Da Re, Olivares, Smith, & Vallejo-Marín, 2020; Grant, 1924; Stace, 2010; Tokarska-Guzik & Dajdok, 2010; Vallejo-Marin & Lye, 2013). Unlike many invasive and non-native species, detailed historic botanical records (Sims, 1812) and travel diaries of early explorers (von Langsdorff, 1817) allow us to clearly retrace the history of the first introduction of *M. guttatus* into Europe. Historical records of M. guttatus reaching the UK paint a clear picture, but beyond this little us known. Here we test the hypothesis that the UK acted as a bridgehead for worldwide invasion.

The first European record of *M. guttatus* appears in *Curtis’ s Botanical Magazine* (Sims, 1812), which presents a plate of *Langsdorff’ s Mimulus* (*Mimulus langsdorfii* Donn ex Sims), featuring a flowering individual of *M. guttatus*. The provenance of the depicted material is from Grigori von Langsdorff who “*brought it, as we are informed, from Unalashka, one of the Fox Islands*” (Unalaska, Aleutian Islands) (Sims, 1812), in his capacity as a naturalist on a Russian expedition to the Alaskan territories in 1805. Langsdorff describes how the expedition reaches Unalaska on 16 July 1805, and, after anchoring in Sea-Otters Bay (probably present-day Ugadaga Bay), they travelled on foot to Iluluk (Dutch Harbor). Here, Langsdorff first encounters *M. guttatus: “splendid flowers were in blow upon the shore, among which a new* Mimulus *and* Potentilla, *which has never yet been described, were particularly to be distinguished*.*”* (von Langsdorff, 1817, p. 329). Material brought by Langsdorff made its way to various Botanic Gardens including Moscow (where is listed as *M. guttatus* Fischer *nom. nudum*) and Montpellier (where De Candolle validly published the name *M. guttatus*). The seeds of *M. guttatus* also reached the Botanic Gardens at Cambridge in 1812, and it is therefore almost certain that the original species description included specimens collected by Langsdorff in Unalaska (Grant, 1924).

Presciently, the *Botanical Magazine* recognized the potential for *M. guttatus* to become established outside western North America, and the 1812 entry states that because the taxon has showy flowers and is “*easily propagated by seeds, and most probably by its runners, must soon be very common*.*”* (Sims, 1812). In fact, the first naturalised populations in the British Isles are recorded by 1830 (Roberts, 1964), rapidly spreading throughout the United Kingdom (UK) (Preston, Pearman, & Dines, 2002). The introduction history of *M. guttatus* outside of the UK is much less well understood. *Mimulus guttatus* seems to have reached New Zealand and become naturalised by 1878 (Owen, 1996), and the introduction of this taxon to eastern North America may have occurred much later in the second half of the 20th century (Murren, Chang, & Dudash, 2009). Therefore, the material brought in by Langsdorff represents the first introduction of *M. guttatus* outside its native range, and the subsequent arrival and naturalisation on the British Isles is the best documented, and currently most widespread, monkeyflower invasion (Da Re et al., 2020; McArthur, 1974; Preston et al., 2002; Roberts, 1964; Stace, 2010; Stace & Crawley, 2015).

The historical hypothesis of an Alaskan origin of European monkeyflowers is consistent with results from previous genetic analysis of *M. guttatus* in the United Kingdom (Pantoja, Simón-Porcar, Puzey, & Vallejo-Marín, 2017; Puzey & Vallejo-Marin, 2014). However, these studies did not include material from the putative origin (Aleutian Islands), and due to their focus on UK populations, did not examine genetic relationships between native populations and introduced populations in other parts of the range such as in Eastern North America, the Faroe Islands, mainland Europe and New Zealand. Native *M. guttatus* presents an enormous breadth of ecological and genetic diversity (Vickery, 1978; Wu et al., 2008), and it remains unknown how much of this diversity is represented among introduced populations and the extent to which non-native populations have diverged. Recently, Da Re *et al*. (2020) used ecological niche modelling to compare the climatic envelope of native and introduced *M. guttatus* populations, finding no evidence of niche shift in the introduced UK populations compared to the native ones. Moreover, the highest niche similarity of invasive UK populations occurred in the Aleutian Islands (Da Re et al., 2020), lending support to the historical hypothesis that traces their origin to Langsdorff.

Here we provide the first global genetic analysis of native and introduced populations of *M. guttatus* by marrying historical information with genomic analyses. Specifically, we: (1) Resolve range-wide relationships at the population level in the introduced range, as well as in the native range including the previously under-sampled regions of the Aleutian Islands and mainland Alaska; and (2) use genomic data to reconstruct the population genetic history of introduced UK populations and test the hypothesis that UK populations have a simple Aleutian origin or are the product of a more complex invasion history.

## Materials and Methods

### Study system and population sampling

*Mimulus guttatus* Fischer ex DC (section *Simiolus*, Phrymaceae), the common monkeyflower, is a widespread species with a native range extending across western North America from northern Mexico to the farthest reaches of the Aleutian Island chain in Alaska (Da Re et al., 2020; Vickery, 1978). The invasive range includes much of the UK, the Faroe Islands, parts of mainland Europe, New Zealand, and Eastern North America (Da Re et al., 2020). The species is self-compatible and predominantly outcrossing (Ritland, 1989). Most populations are diploid, although tetraploid populations occur throughout the native range (Vickery, Crook, Lindsay, Mia, & Tai, 1968) and tetraploid populations have also evolved in the introduced range (Simón-Porcar, Silva, Meeus, Higgins, & Vallejo-Marín, 2017; Vickery et al., 1968). In the native range, populations comprise either small annual plants that reproduce exclusively by seed or perennial plants that reproduce by both seed and vegetative stolons. Only perennial plants are documented in the invasive range.

We sampled populations of *M. guttatus* in the native range of western North America and the main areas of introduction in eastern North America, Europe and New Zealand for a total of 521 individuals from 158 populations (Figure 1, Table 1). In the native range, the samples included 70 previously genotyped populations (Twyford & Friedman, 2015), spanning Arizona to British Columbia, plus an additional population from Vancouver Island. To fill the gap of previous studies, and to specifically address the hypothesis of an Alaskan origin of introduced UK populations, we collected samples from 32 populations in Alaska, including 14 populations from the Aleutian Islands (Attu, Unalaska, Akutan and Unimak) (Table S1). Voucher specimens of the newly sampled populations are deposited in the University of Alaska herbarium (ALA). In the introduced range, we sampled four populations in eastern North America, one from the Faroe Islands, one from Germany, six from New Zealand, and 43 from UK populations from Cornwall to the Shetland Islands. As an outgroup we included three diploid individuals from a population of *M. glabratus* from Michigan, USA. We also sampled three tetraploid UK *M. guttatus*, 19 individuals of *M. luteus* from both native and introduced ranges (with which *M. guttatus* hybridises in the introduced range to produce a sterile but widespread triploid, *M. x robertsii*), three *M*. × *robertsii*, and three *M. peregrinus* (the allohexaploid species derived by whole genome duplication from *M. x robertsii*; (Vallejo-Marin, Buggs, Cooley, & Puzey, 2015) (Table S1). In total, we had samples from 103 populations of *M. guttatus* from the native range, and 55 populations from the introduced range (Table 1). Full sample details are provided in Table S1.

**Table 1.** Summary of the number of populations and individuals sampled and sequenced. A detailed breakdown by population is shown in Table S1.

|  | Region | Number of<br>populations | Number of<br>individuals |
| --- | --- | --- | --- |
| Native | Western North America<br>(excluding Alaska) | 71 | 182 |
|  | Western North America<br>(Alaska only) | 32 | 106 |
| Introduced | Eastern North America | 4 | 34 |
|  | Faroe Islands | 1 | 4 |
|  | United Kingdom | 43 | 161 |
|  | Germany | 1 | 9 |
|  | New Zealand | 6 | 25 |
| Total |  | 158 | 521 |

**Figure 1.**
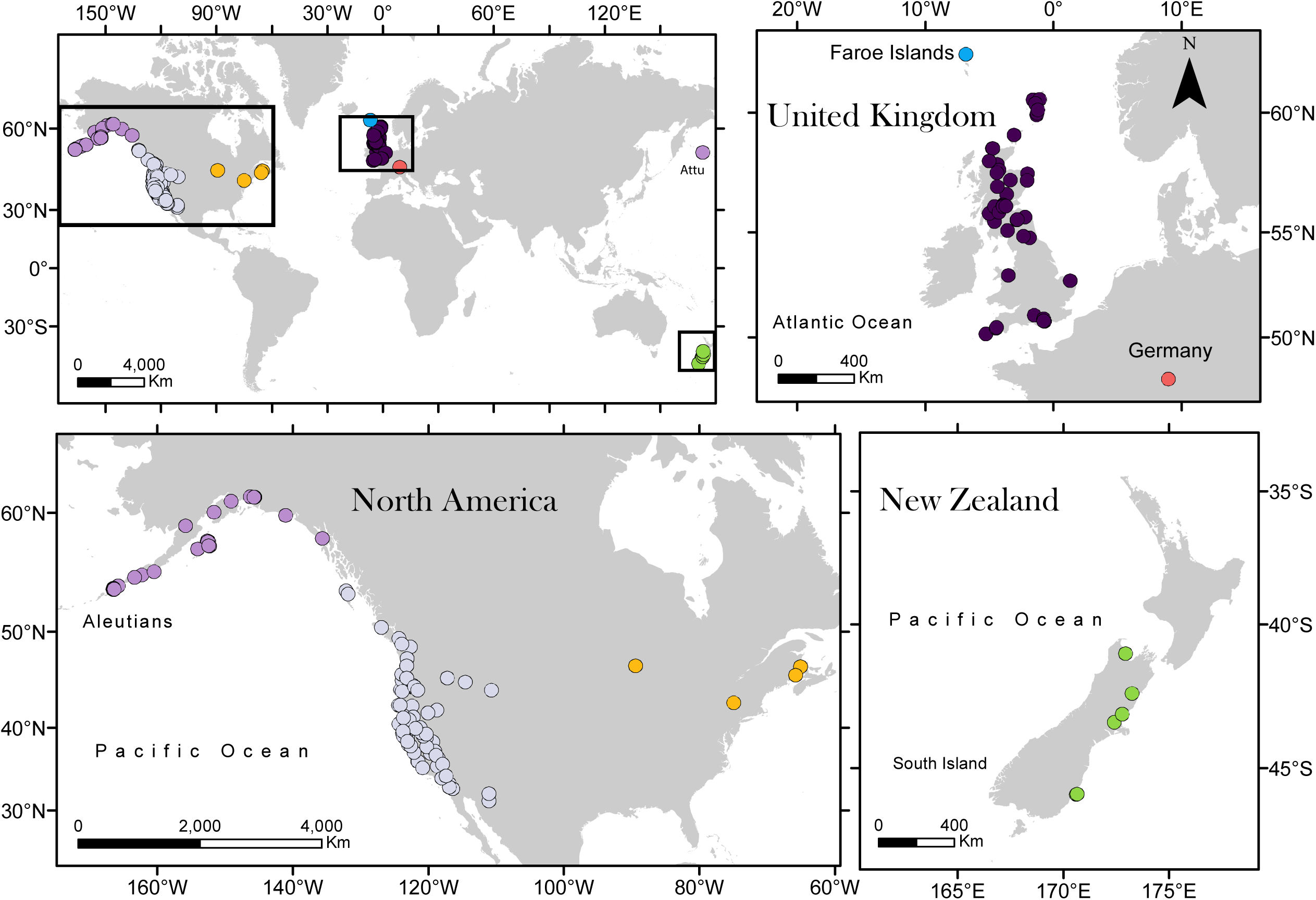
Global sampling of *Mimulus guttatus* populations. Native populations in western North America are shown in green in the inset.

### Genotyping

To obtain DNA for genotyping, we germinated field-collected seeds from all new populations in a controlled environment facility at the University of Stirling. We extracted genomic DNA from fresh leaves or flower buds using the DNeasy Plant Kit (Qiagen, Germantown, MD), with samples standardised to 100ng DNA for library preparation. We used genotyping by sequencing (GBS) to generate genome-wide polymorphism data (Elshire et al., 2011). For GBS library preparation, we used the same protocol as Twyford and Friedman (2015), using the enzyme PstI, and pooling samples in a 95-plex (plus one blank water control) for 100bp single-end sequencing on the Illumina HiSeq 4000 at the University of Oregon. We analysed raw sequence reads using the Tassel5-GBSv2Pipeline (Glaubitz et al., 2014), using the *M. guttatus* v2 genome (Hellsten et al., 2013) as a reference. For population genetic analyses, we retained only variable sites (SNPs), but for tree reconstruction, we generated a sequence matrix with both SNPs and invariant sites (setting MAF = 0).

### Tree building

We sought to resolve evolutionary relationships between populations and species using polymorphism-aware phylogenetic models implemented in IQ-TREE (Nguyen, Schmidt, von Haeseler, & Minh, 2015). These models use population site frequency data, and therefore account for incomplete lineage sorting (Schrempf, Minh, De Maio, von Haeseler, & Kosiol, 2016). This phylogeographic approach generates an initial visualisation of population history and broad scale geographic genetic structure from the genome-wide signal, prior to more detailed characterisation with population-level approaches (described below). We analysed two datasets, one for all sampled *Mimulus* taxa, and one for *M. guttatus*, with both datasets including *M. glabratus* as an outgroup. Each analysis used the full GBS sequences with invariant sites, filtered to include 8,798 sites with less than 50% missing data. We calculated population allele frequencies using the counts file library (cflib) python scripts that accompany (Schrempf et al., 2016). Model-fitting was performed with ModelFinder (Kalyaanamoorthy, Minh, Wong, von Haeseler, & Jermiin, 2017). IQ-TREE analyses subsequently used the best-fitting model (TVM+F+G4) allowing for excess polymorphism (+P) and with five chromosome sets per population (+N5). Tree searches were performed with settings recommended for short sequences, including a small perturbation strength (-pers 0.2) and large number of stop iterations (-nstop 500). Topological support was assessed using an ultrafast bootstrap approximation approach (Minh, Nguyen, & von Haeseler, 2013), with 1000 bootstrap replicates. Trees were visualised with *FigTree* (Rambaut, 2014).

### Population genetic structure

For population genetic analyses in *M. guttatus*, we filtered the SNP data (44,552 loci from 521 *M. guttatus* individuals) using VCF Tools and kept only biallelic loci that were genotyped in at least 75% of all individuals, which reduced the number of genotyped SNPs to 1,820 loci. We then removed individuals with less than 50% genotyped loci, reducing the number of individuals from 521 to 474. Finally, we used PLINK to thin the data set to reduce linkage disequilibrium among SNPS using a pairwise correlation coefficient of 0.5 (--*indep-pairwise 50 5 0*.*5*). The final *M. guttatus* dataset contained 1,498 SNPs from 474 individuals in 155 populations.

To analyse population genetic structure, we conducted a principal component analysis using the *glPca* function in *adegenet* (Jombart & Ahmed, 2011) in *R* ver. 4.0.0 (R Development Core Team, 2020). We used *K*-means grouping implemented with the function *find*.*clusters* in *adegenet* to identify clusters of individuals in the data without using a priori groupings. For this analysis, we used 100 randomly chosen centroids for each run, and calculated the goodness of fit for each model for values of *K* between two and 15. For the selected *K* value, we also ran a Discriminant Analysis of Principal Components (DAPC) (Jombart, Devillard, & Balloux, 2010) using the inferred groups for assigning individual membership. We further used *fastStructure* (Raj, Stephens, & Pritchard, 2014) to infer population structure across *M. guttatus* populations using a Bayesian framework. For this analysis, we randomly subsampled the data to include a maximum of three individuals per population (408 individuals in total) from both native and introduced ranges, and analysed values of *K* from 2-8.

### Introduction history reconstruction by ABC

Our preliminary analyses indicated that introduced *M. guttatus* had a complex origin with multiple introductions in different non-native regions. In order to gain a more detailed understanding of the demographic history of non-native populations, we focused on the introduction of *M. guttatus* to the UK, which has been best studied both historically and genetically (Pantoja et al., 2017; Puzey & Vallejo-Marin, 2014). Therefore, we implemented an approximate Bayesian computation (ABC) approach to determine the most likely *M. guttatus* introduction history in the UK. For this analysis, we used the pruned data set consisting of 1,498 SNPs but included only individuals from the native range or the UK (399 individuals). Individuals from the native range were grouped into one of five groups (“genetic group”) delimited by the genetic clustering and phylogenetic tree analysis (see Results section): North (NORTH; N=62), South (SOUTH; N=42), Coastal (COAST; N=30), Alaska and British Columbia (AKBC; N=70) or Aleutian (ALE; N=45). Six individuals from two populations (SWC and HAM) that formed a separate genetic group in the native range were not included in this analysis. Individuals from the UK were considered to belong to a single population (UK; N=150).

Because all possible scenarios of divergence between the five native groups would have been computationally impossible to test, native group genetic relationships were determined from the phylogenetic tree topology (see Results section). All the simulations assumed that the North population diverged from an ancestral population at time t_4_, from which the South population diverged at time t_5_. In addition, the Coastal population diverged from the ancestral population at time t_3_ from which the Alaska-British Columbia population diverged at time t_2_, and the Aleutian population diverged from there at time t_1_. The simulated demographic models share this native population divergence history and only differed by their introduction history into the UK.

We first considered simple introduction models where the UK population was derived from a single native origin at time t_0a_ (models A1 to A5, Supporting Materials File 1). We then simulated UK introduction from a single origin at time t_0a_ followed by a second introduction at time t_0b_ (two-waves introduction models; models B). This strategy resulted in the definition of eight different two-waves introduction models (models B1 to B8, Supporting Materials File 1). We then tested more complex introduction models using a similar logic, modelling three-waves (models C1 to C9), four-waves (models D1 to D8) and five-waves (models E1 to E5) introduction models by integrating the most likely origins identified in previous sets of models to define a restricted number of models to compare. A full version of other assumptions and simulation parameters is given in Supplemental Materials S1.

For each demographic model, we simulated 10,000 genetic datasets consisting of 1435 independent SNP genotypes for 798 haploid individuals distributed following the sample size of all six populations in the real dataset using *Fastsimcoal2* version 2.6.0.3 (Excoffier, Dupanloup, Huerta-Sanchez, Sousa, & Foll, 2013) called by *ABCtoolbox* version 1 (Wegmann, Leuenberger, Neuenschwander, & Excoffier, 2010). We passed a custom bash script to *ABCtoolbox* to add missing genotypes to the simulated dataset at an identical rate to the observed level in the real data. Then, we used *ABCtoolbox* to call the *arlsumstat* program (Excoffier & Lischer, 2010) to compute summary statistics from the simulated genotypes. We computed all available statistics within and between populations for bi-allelic loci (67 summary statistics). In addition, we computed summary statistics within and between three defined regional groups (NORTH and SOUTH in one group; COASTAL, AKBC and ALE in a second group; and UK in a third group) representing an additional set of 29 summary statistics.

### ABC model comparisons

We performed iterative model comparisons by comparing increasingly complex models (Table 2). In the first round, the introduction models assume a single introduction from one of the five native genetic groups. Then in round two, we considered two introductions models that necessarily involved the population origin from round one. This allowed us to define two sets of two-waves introduction models: One set consisting of four models with the most likely origin in previous rounds as the first introduction origin, followed by a second introduction from one of the four other native populations. And a second set of four models, which assume that the most likely origin in the previous round constitutes the second introduction, while the first introduction originated from one of the four other native populations (Table 2). We compared the most likely single introduction model and the eight two-waves introduction models. We then considered more complex models, comparing nine three-waves introduction models and the most likely single and two-waves introduction models (Table 2). We subsequently compared models assuming four-waves and five-waves of introduction while still including more simple models in the comparisons (Table 2). Demographic models were compared using a random forest approach implemented in the *R* package *abcrf* (Pudlo et al., 2016).

**Table 2.**
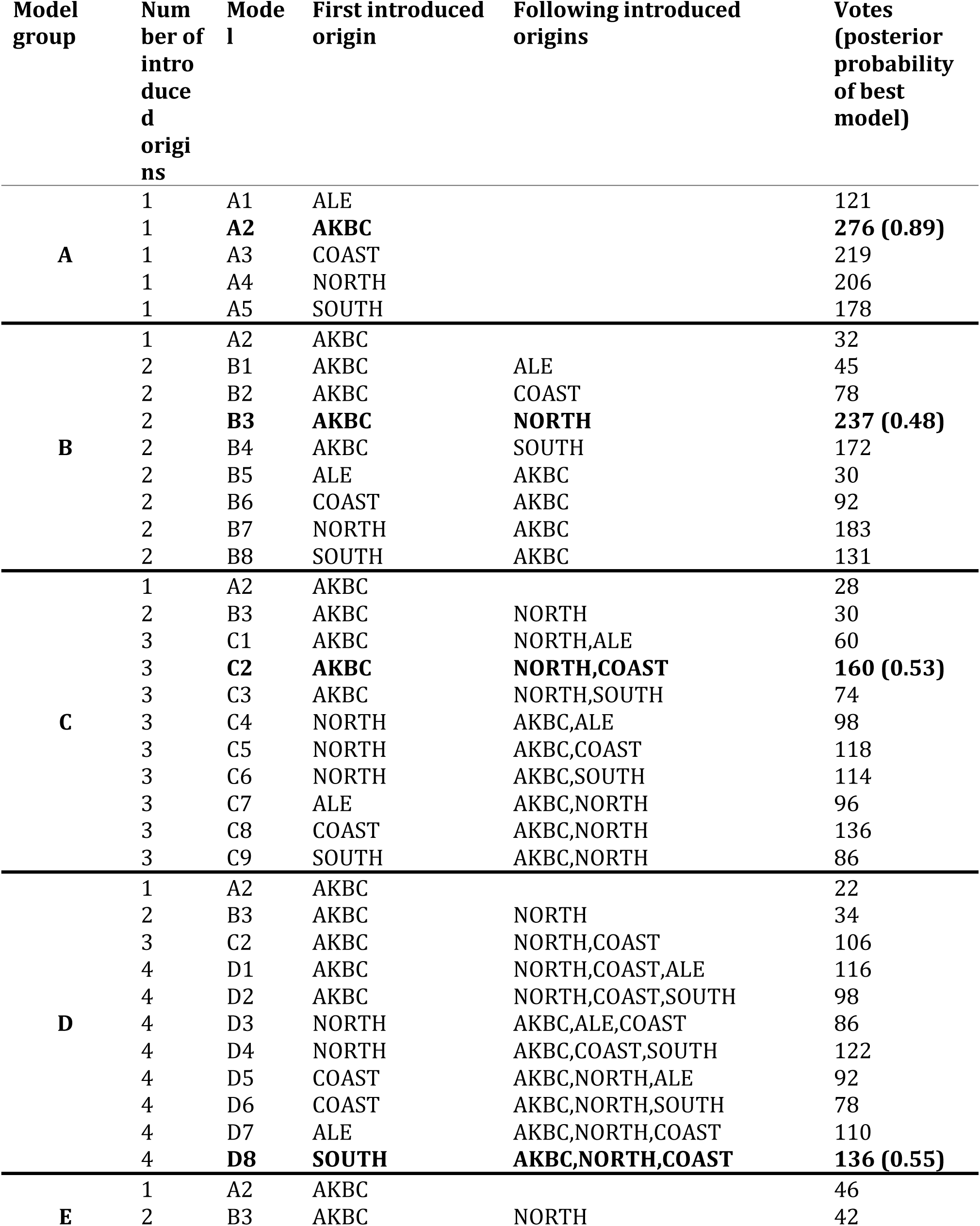
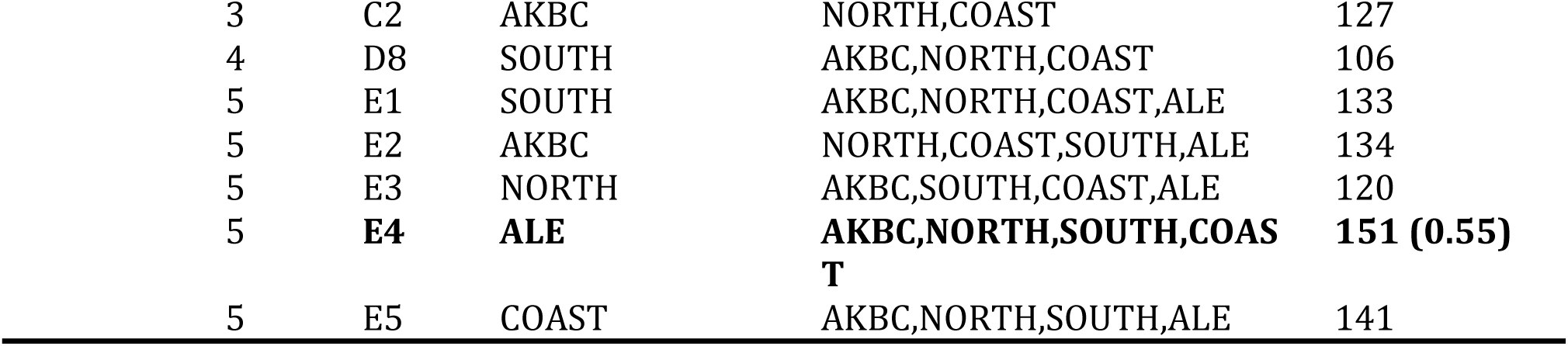
Stepwise comparison of demographic models of the invasion of *Mimulus guttatus* into the United Kingdom using 10,000 simulations for each of the model and random forest ABC model selection approach. At each step (model groups A-E), more complex introduction histories are considered while keeping the most likely models selected in previous comparison steps. The most likely model at each step is indicated in bold.

We built a classification random forest model using 1000 trees and a training dataset consisting of the summary statistics computed for the 10,000 simulated genetic datasets for each model. We estimated the classification error rate for each model using an “out-of-bag” procedure to quantify the power of the genetic data given the models and prior distribution specifications to differentiate the different demographic models. Then, we used the summary statistics computed based on the observed genotypic data to predict the demographic model that best fit the data using a regression forest with 1000 trees. We report the number of “votes” for each demographic scenario and the approximation of the posterior probability of the most likely model. We used the overall most likely scenario to simulate 100,000 genetic datasets using parameters and prior distributions described above to estimate demographic model parameters. We built a regression random forest model implemented in *abcrf* based on the summary statistics using 1000 trees. We estimated the posterior median, 0.05 and 0.95 quantiles of the model parameters by random forest regression model based on the summary statistics of the observed genotypic composition.

## Results

### Demographic relationships in the native range

The global sampling of *M. guttatus*, including populations sampled across ∼5000km of its distribution in North America (Figure 1), allowed us to resolve demographic groupings in both native and introduced ranges. In the native range, including the newly sampled Alaskan region, strong geographic structure is evident from phylogenetic analysis (Figure 2), with four well-resolved North, South, Coastal and North Pacific clades (Twyford & Friedman, 2015). The newly sampled populations in Alaska and the Aleutian Islands form part of the North Pacific Clade (Figure 2). This clade is sister to the Coastal clade and includes populations from northern Washington to the westernmost Aleutian Islands (Attu Island). Phylogenetic analysis revealed an unexpected placement of some populations from inland Oregon, including those from Iron Mountain, which conflicts with previous analyses and their expected relationships based on simple geography. The tetraploid *M. guttatus* population sampled in the Shetland Islands in the UK is nested among other geographically proximate populations, further supporting the local origin of this autopolyploid in the introduced range (Simón-Porcar et al., 2017). Finally, *M. luteus* formed a strongly supported clade, and the triploid and allohexaploid hybrids, *M x robertsii and M. peregrinus* can be clearly distinguished from both parental taxa (*M. guttatus* and *M. luteus*).

**Figure 2.**
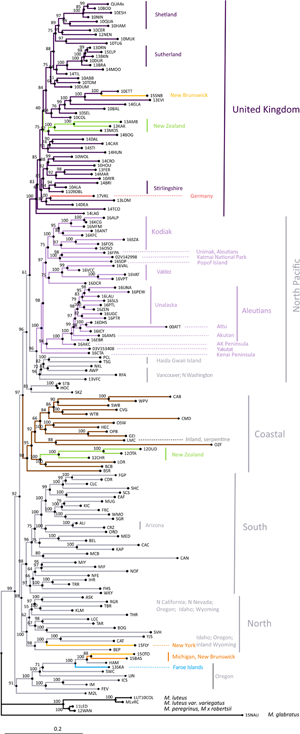
Maximum likelihood phylogenetic reconstruction of the relationship between studied *Mimulus guttatus* populations, and including populations from *M. luteus* (LUT10COL, UK), *M. luteus var. variegatus* (MLvRC, Chile) *M x robertsii* (12WAN) and *M. peregrinus* (11LED). The tree is rooted using a population of *M. glabratus* from Michigan (15NAU)

### Global invasion of *Mimulus guttatus*

At a global scale (Figure 1), introduced *M. guttatus* populations are scattered across the phylogeny, indicating many independent introductions from across the native range (Figure 2). In contrast, however, all UK *M. guttatus* populations form a sister group to the North Pacific clade. The UK group also includes other non-native populations from New Zealand, Canada and Germany, suggesting it may be the source for these. Other New Zealand populations are grouped within the Coastal clade, suggesting a potential second introduction. Moreover, interesting geographic discontinuities exist in North America, with a non-native New York population nested in the native North clade. Finally, two additional populations from eastern North America, as well as the single sampled population from the Faroe Islands are grouped together with the native HAM-SWC group from Oregon (Figure 2). Thus, the UK populations are genetically similar to each other and are closely related to some of the introduced populations of *M. guttatus* in New Zealand and eastern North America. However, the placement of other non-native populations within various native clades clearly indicates additional, independent introductions to New Zealand, eastern North America and the Faroe Islands, suggesting a complex history of colonisation.

Among native populations those from the UK form a separate genetic cluster, as seen in principal component analysis (PCA) (Figure 3). As in the phylogenetic reconstruction, the UK group is closely associated with non-native populations from New Zealand, Germany and eastern North America. The PCA is also consistent with two separate introductions into New Zealand, one of them closely related to UK populations, and three independent origins of non-native populations in eastern North America. One of these origins of eastern North American populations is shared with the population from the Faroe Islands, forming a distinct group with two native populations from Oregon (SWC and HAM; Figure 3). An interactive version of Figure 3 with labelled individuals and populations is available at https://plot.ly/~mvallejo6/1/. Population structure in the native range is less clear from the worldwide PCA, although the North Pacific clade and particularly the Aleutian Islands populations are well differentiated along the first principal component (Figure 3).

**Figure 3.**
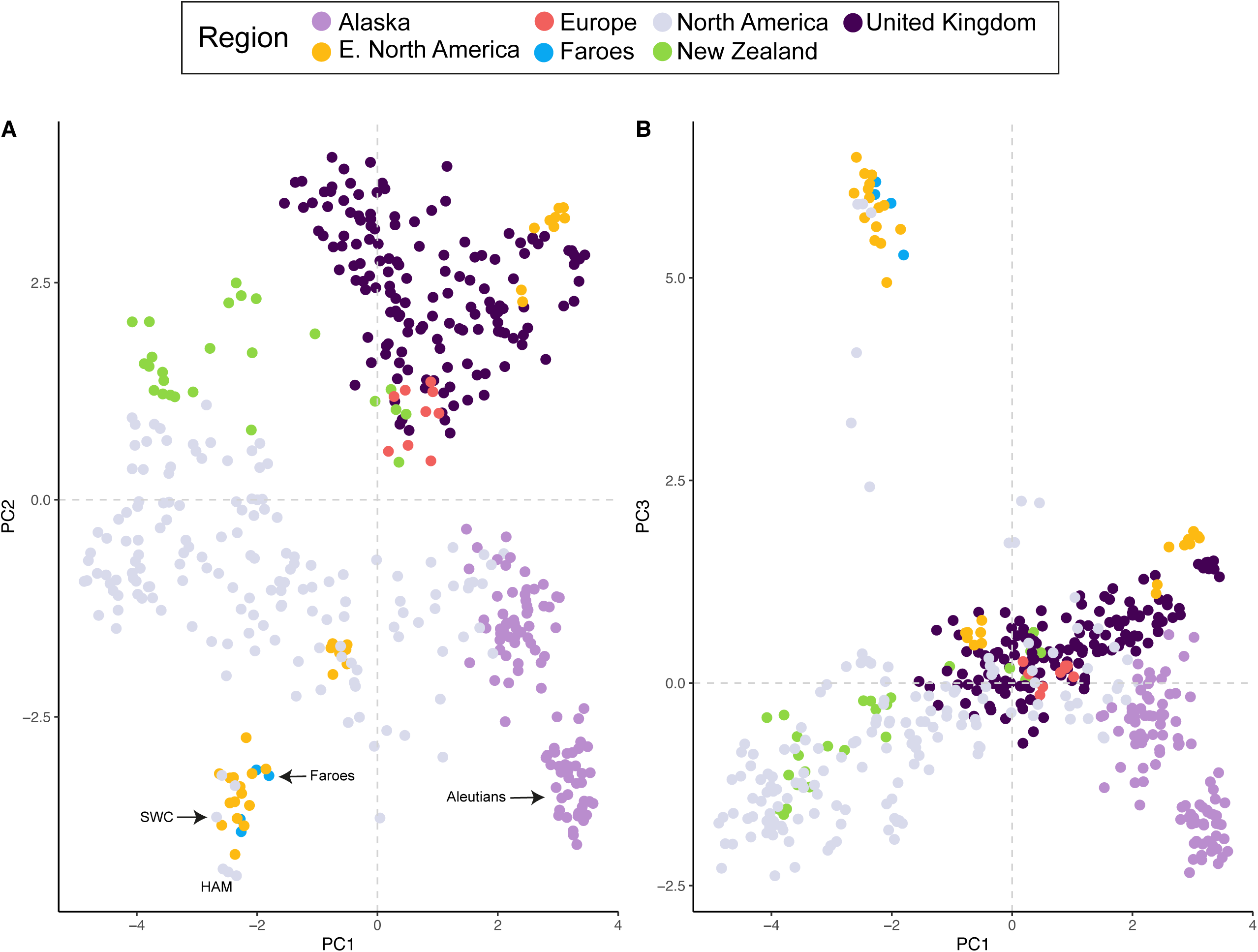
Principal Component Analysis (PCA) of 474 individuals of *Mimulus guttatus* from both native and introduced populations genotyped at 1,498 binary SNP loci. (A) Scatterplot of the first two principal components (PC2 vs PC1). (B) Scatterplot of first and third principal components (PC3 vs PC1). Colours indicate sample regions. An interactive 3D figure with individually labelled data points is available at: https://plot.ly/~mvallejo6/1/

Worldwide groupings by *K*-means cluster analysis (Figure 4) partition North American samples are into three groups, New Zealand into two groups, and the single populations from the Faroe Islands and Germany in one group each, largely consistent with the results above. Non-native UK populations form two groups, one mixed with European and Eastern North American samples, and another with New Zealand samples. Native, non-Alaskan populations are distributed in five groups. Aleutian populations form a separate group not shared with other geographic regions. The *fastStructure* analysis with the selected *K* =8 value (Figure S2) provides further support for these groupings. UK populations form a separate group with which multiple affinities with New Zealand and eastern North American samples are evident. Furthermore, the distinctiveness of Aleutian populations relative to other native populations is also obvious (e.g., cluster 4 at K = 8, Figure S2).

**Figure 4.**
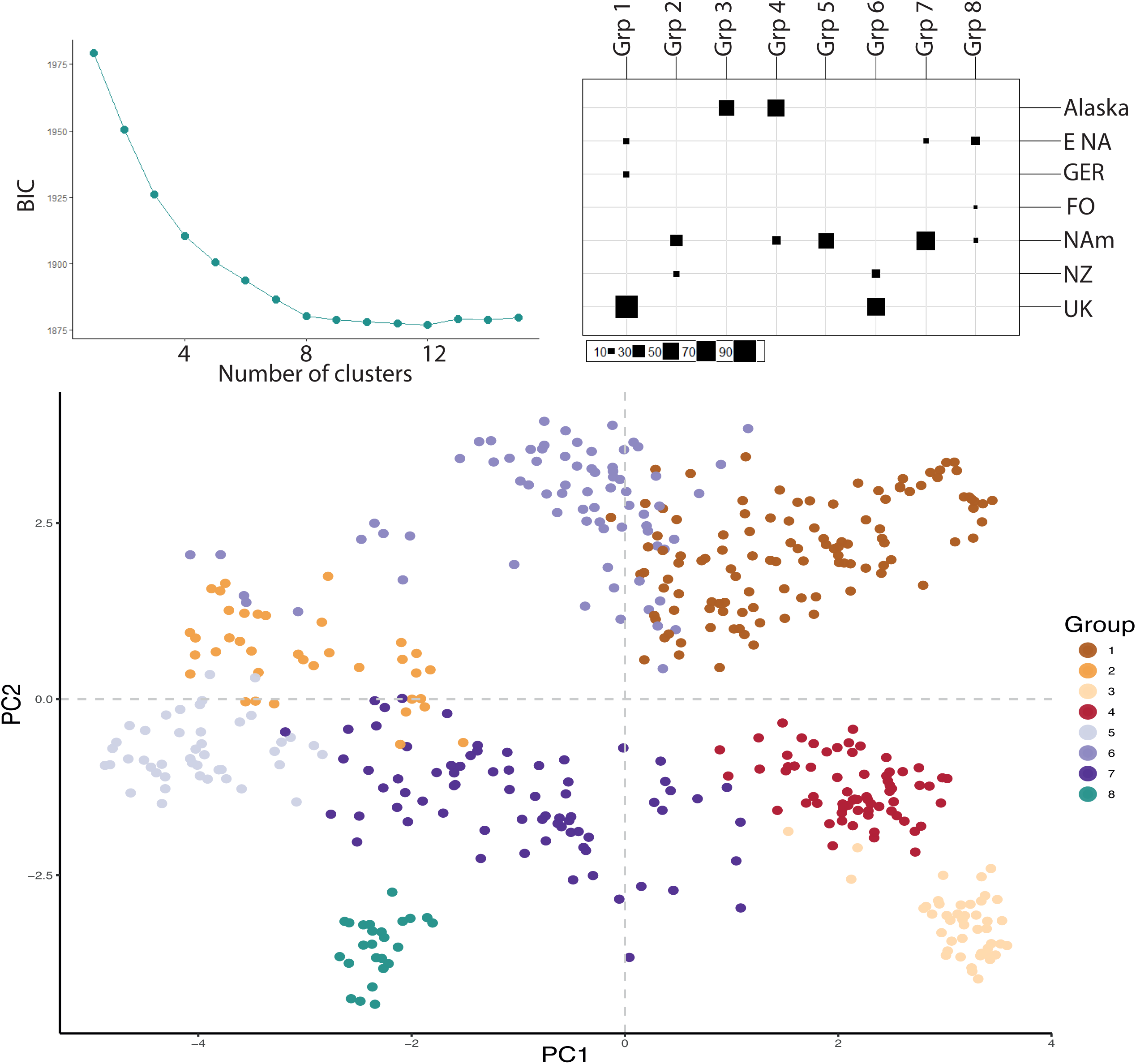
K-means clustering analysis of native and introduced populations of *Mimulus guttatus*. The analysis is based on the first 300 Principal Components. (**A**) Bayesian Information Criterion values for models ranging from 2 to 15 clusters. (**B**) Group membership of each geographic group for the optimal number of clusters (K=8). (**C**) Principal Component Analysis depicted in Figure 3 but coloured by the groups identified in the K-means cluster analysis (K=8). Colours indicate sample regions as follows: Alaska = Alaska; E NA = Eastern North America; GER = Europe (Germany); FO = Faroe Islands; NAm = Western North America; NZ = New Zealand; UK = United Kingdom.

### Introduction history in the UK

To estimate a most likely scenario for the origin and history of introduction of UK populations, we next performed a coalescent analysis with ABC. Our analysis of demographic models allowed us to compare different scenarios for the origin and history of introduction of UK populations relative to five genetic groups in the native range: Aleutians (ALE) and Alaska-British Columbia (AKBC), both of which form part of the North Pacific clade, and the North (NORTH), South (SOUTH), and Coastal (COAST) clades (see Figure 2). When assuming a single introduction event, the most likely source of UK individuals is the AKBC group (Table 2, posterior probability *p*=0.89). However, model comparisons favour scenarios with additional waves of introductions (Table 2). When we model two introductions, a first introduction from AKBC followed by a second introduction wave from NORTH has greatest support (Table 2, *p*=0.48) and is more likely than a single introduction scenario (237 votes against 32 votes, Table 2). Similarly, three introduction models result in selecting an introduction history with a first introduction from AKBC followed by additional introductions from NORTH and COAST (*p*=0.53, Table 2) and then four introduction models identify a first introduction from SOUTH followed by additional introductions from AKBC, NORTH and COAST as the most likely scenario (*p*=0.55, Table 2). Finally, when comparing all best one- to four-wave introduction models, with all possible five-wave introduction models, the most likely introduction history identified consisted of a first introduction from ALE followed by four subsequent waves from the AKBC, NORTH, SOUTH and COAST (E4 model; *p*=0.55, Table 2). Full demographic parameters (e.g., estimated population sizes and introduction times per genetic group; E4 model) are presented in Table S2.

Classification of the datasets simulated under a five-wave introduction scenario showed that 83.4% of the simulations classified were correctly assigned to a five-wave introduction scenario, and 23.7% to the correct model (E4) (Table 3). Thus, the combination of the type and number of molecular markers and model prior specifications we used here contain enough information to confidently differentiate scenarios with different number of introductions (e.g., single introduction vs five-wave introductions). Nevertheless, distinguishing the most likely scenario among these complex and sometimes very similar five-wave introduction scenarios proved more difficult (Table 3, Supporting Material File 1). In other words, our ability to distinguish the order of introductions of the five genetic groups is more limited.

**Table 3.** (**A**) Power to discriminate between alternative demographic models using an “out-of-bag” procedure given the parameter model specification. The comparisons are made at the final selection step between the most likely one- to four-wave introduction models and all possible five-wave introduction models. The table shows how many of the 10,000 simulated datasets generated under a given scenario (A2 to E5, rows) were classified into each demographic scenario (A2 to E5 columns). The number of incorrect classifications is then used to compute the overall classification error. The last column shows the percentage of simulated models classified as E4 (which was the most likely scenario for the observed genetic dataset). Bold numbers indicate correct classification, and underlined numbers indicate >10% incorrect classification. (**B**) Probability of a given number of origins given that the E4 model is selected.

**A.**
| Classified models | A2 | B3 | C2 | D8 | E1 | E2 | E3 | E4 | E5 | Total | Classification error | Probability that E4 is selected |
| --- | --- | --- | --- | --- | --- | --- | --- | --- | --- | --- | --- | --- |
| Simulated models |  |  |  |  |  |  |  |  |  |  |  |  |
| A2 | **8902** | <u>1043</u> | 24 | 12 | 2 | 9 | 0 | 1 | 7 | 10000 | 11.0% | 0.0% |
| B3 | <u>2151</u> | **7616** | 26 | 15 | 4 | 73 | 11 | 25 | 79 | 10000 | 23.8% | 0.3% |
| C2 | 210 | 463 | **4844** | <u>1576</u> | 675 | 642 | 534 | 469 | 587 | 10000 | 51.6% | 5.5% |
| D8 | 447 | 330 | <u>3067</u> | **1905** | <u>1039</u> | 662 | 908 | 911 | 731 | 10000 | 81.0% | 10.7% |
| E1 | 355 | 336 | <u>2094</u> | <u>1342</u> | **1462** | 735 | <u>1323</u> | <u>1339</u> | <u>1014</u> | 10000 | 85.4% | 15.8% |
| E2 | 400 | <u>1317</u> | <u>2161</u> | <u>1039</u> | 643 | **1473** | 756 | <u>1062</u> | <u>1149</u> | 10000 | 85.3% | 12.5% |
| E3 | 28 | 625 | <u>1894</u> | <u>1173</u> | <u>1277</u> | 857 | **1641** | <u>1438</u> | <u>1067</u> | 10000 | 83.6% | 17.0% |
| E4 | 344 | <u>1096</u> | <u>1024</u> | 993 | <u>1219</u> | 918 | <u>1313</u> | **2009** | <u>1084</u> | 10000 | 79.9% | **23.7%** |
| E5 | 435 | <u>1291</u> | <u>1491</u> | 777 | 963 | <u>1170</u> | <u>1006</u> | <u>1228</u> | **1639** | 10000 | 83.6% | 14.5%<sup>739</sup> |
| **Total** | **13272** | **14117** | **16625** | **8832** | **7284** | **6539** | **7492** | **8482** | **7357** |  |  | 740 |

**B.**
| Number of origins of UK populations | Probability given that the E4 scenario is selected |
| --- | --- |
| 1 | 0.0% |
| 2 | 0.3% |
| 3 | 5.5% |
| 4 | 10.7% |
| **5** | **83.4%** |

The posterior probability of 55.1% for the E4 model (Table 2), supports a first introduction from ALE followed by additional introductions from the other four other origins (Figure S3). However, most of the posterior distributions of demographic parameters (e.g., effective population size, number of generations since introduction) for model E4 were nearly identical to the prior distributions (Table S2, Supporting Material File 2), indicating limited information content of the genetic dataset to estimate the demographic parameters of this complex introduction history.

## Discussion

Here we provide the first global picture of the genetic relationships between native and introduced populations of *Mimulus guttatus*, including targeted sampling of a historically-indicated origin for the UK bridgehead population. Our results can be summarised in three main findings: (1) *Mimulus guttatus* achieved a broad distribution across geographic boundaries through multiple repeated introductions from genetically distinct source populations; (2) In some cases, the establishment of *M. guttatus* in the invasive range was achieved via a bridgehead process, where invasive populations serve themselves as sources for further, more distant vanguard invasions. This is well illustrated in our discovery of the establishment of invasive populations in New Zealand and eastern North America by way of UK invasive populations; (3) Admixture in the introduced range has given rise to genetically distinct populations generating novel genetic, and therefore phenotypic, combinations.

### Multiple introductions and bridgehead invasions

Widely distributed taxa that serve as a source of invasive populations pose a particular challenge for molecular studies aiming to reconstruct the history of biological invasions. The distribution of *M. guttatus* spans from Mexico to the Aleutians and covers more than 6000km of coastline (Vickery, 1978). To identify potential sources of specific invasion events, sampling large geographic regions is required. *Mimulus guttatus* has been the subject of continuous study for the last 60 years (Wu et al., 2008), and previous work has collected population samples across nearly its entire native range (Friedman, Twyford, Willis, & Blackman, 2015; Lowry, Hall, Salt, & Willis, 2009; Oneal, Lowry, Wright, Zhu, & Willis, 2014). Our analyses of large-scale population samples from the native range builds on the recent finding of geographic genetic structure corresponding to separate coastal and northern colonisation events in North America (Twyford et al., *in press*). Here we fill-in crucial gaps with sampling from Alaska and the Aleutian Islands, which reveals strong geographic structure in the far north west of the species range, with genetic clusters by islands in the Aleutians. This extensive sampling in the native range allows us to show that Aleutian populations have acted as important conduits to the invasion of *Mimulus* in Europe and beyond.

Many biological invasions by both plants and animals are associated with multiple introductions, to the extent that single introduction invasions are considered the exception (Dlugosch & Parker, 2008). Here we found clear evidence that introduction of *M. guttatus* into various geographic regions has occurred by colonisation from multiple genetically distinct sources. For example, among the four populations we sampled in eastern North America, where *M. guttatus* was introduced in the last century, there is evidence of three genetically distinct groups, one of which also occurs in the Faroe Islands (Figure 3). Similarly, introduced populations in New Zealand have at least two separate genetic origins, including a close affinity with native populations (near Santa Cruz, California) located 11,000km away and with non-native populations in the UK. The multiple origins of invasive populations found in the same geographic region is important for several reasons. From a management perspective, multiple introductions can help identify locations of transport routes that are susceptible for further invasions. Moreover, multiple introductions may help invasive populations overcome demographic and genetic bottlenecks associated with introduction events (Dlugosch & Parker, 2008). In species that are introduced via the ornamental trade, as was probably the case for monkeyflowers, repeated introductions may not be unusual. To date it is still possible to freely purchase monkeyflowers in UK garden centres. However, because the type sold is no longer *M. guttatus* but horticultural varieties of its close relative *M. luteus*, we speculate that the multiple introductions detected in the invasive range of *M. guttatus* reflect historical events (19_th_ and 20_th_ centuries) rather than recent reintroductions. In addition, we did not find evidence of large-scale admixture from *M. luteus* shaping genetic variation in *M. guttatus*, consistent with the strong reproductive barriers imposed by differences in ploidy level between these *Mimulus* taxa (Meeus, Šemberová, De Storme, Geelen, & Vallejo-Marín, 2020).

The genetic history of these invasions reveals a complex series of introduction events associated with early establishment (19_th_ century). Our ABC analyses reconstruct this history and show that extant populations are composed of a combination of multiple genetic groups from across the native range. Reconstruction of demographic events during introduction (Figure 7) supports an initial introduction of *M. guttatus* from the Aleutian Islands, which is consistent with the historical records of Langsdorff’ s expedition and subsequent transfer of material to Russian, European and British collections. The colonisation of the UK by these exotic Aleutian monkeyflowers may have been facilitated by the close similarity of the ecological niche of *M. guttatus* in the British Isles and the Aleutian Islands (Da Re et al., 2020). Climatic pre-adaptation of Aleutian monkeyflowers provided early arrivals with an opportunity for initial establishment. It is also clear that an initial introduction from the Aleutian Islands was accompanied or quickly followed by multiple introductions from other parts of the range. The UK seems to have become a melting pot for *M. guttatus* resulting in admixture of previously differentiated populations, which resulted in the creation of a unique set of genotypes that are now characteristic of UK populations (Figures 4 & 5).

Invasive populations can themselves become sources for subsequent invasions, a phenomenon termed the “bridgehead effect” (Lombaert et al., 2010). For example, the invasion of Australia by ragweed (*Ambrosia artemisiifolia*, Asteraceae) occurred not from native North American populations, but from populations in the introduced European range (van Boheemen et al., 2017). Our results indicate that UK populations served as a stepping-stone for secondary invasions in other parts of the non-native range. This bridgehead effect in invasive monkeyflowers is most clearly illustrated in the invasion of New Zealand. Some invasive populations there share a close genetic affinity to UK populations. The genetic similarity is consistent with the exchange of biological material, including horticultural taxa, in the 19_th_ century, as British people migrated to New Zealand (Bridge & Fedorowich, 2004). The single sampled population in continental Europe (Germany) also shows a close relationship to UK populations. Unfortunately, without further sampling it is difficult to establish whether UK populations contribute to the extant populations of *M. guttatus* in Europe. Morphologically, *M. guttatus* populations in Russia, Germany and the Czech Republic resemble UK material (Vallejo-Marín, *pers. obs*.) but the genetic identity of continental Europe populations remains to be investigated. In this regard, genomic analyses of herbarium specimens could provide important additional insights (Gutaker, Reiter, Furtwangler, Schuenemann, & Burbano, 2017). Particularly tantalising would be to compare specimens from herbaria in Russia, France and the UK, where historical links connect early *Mimulus* collections with Langsdorff’ s expedition to Alaska in the early 19th century. Finally, we also detected a close affinity between UK populations and a population in the non-native range in eastern North America. Populations of *M. guttatus* in eastern North America are generally small, occurring in the states of Michigan, New York, USA and in New Brunswick, Canada (Murren et al., 2009). These small and sparsely distributed populations show diverse genetic origins and seem to be much more recently established (second half of the 20_th_ century). The mechanism of introduction of UK material into eastern North America is unknown but it could be associated with horticultural exchanges (Chapman et al., 2017; Haeuser et al., 2018; Seebens et al., 2015).

### Admixture and adaptive potential

Multiple introductions and admixture can, in principle, both increase or decrease the performance and adaptive potential of invasive populations (Barker et al., 2019; Rius & Darling, 2014; Verhoeven, Macel, Wolfe, & Biere, 2011). Multiple introductions from genetically distinct sources introduce variation and alleviate the negative effects of demographic bottlenecks associated with colonisation. Moreover, genetically diverse populations are less likely to experience the deleterious effects of inbreeding depression (Dudash, Murren, & Carr, 2005; Verhoeven et al., 2011) and can increase individual fitness through heterosis (Rius & Darling, 2014). In contrast, admixture may reduce overall fitness if gene flow results in outbreeding depression (Frankham et al., 2011), a phenomenon that can occur due to epistatic interactions or, for example, the breakdown of locally adapted genotypes. In *M. guttatus*, experimental work indicates that both positive and negative effects of admixture can be observed in invasive populations. For example, crossing native and introduced populations results in an increase in biomass, and both clonal and sexual reproduction in greenhouse conditions (Li, Stift, & van Kleunen, 2018; van Kleunen, Rockle, & Stift, 2015). In field conditions, the effects of admixture can be reversed, and a common garden study shows that admixture between UK *M. guttatus* and both annual and perennial populations from the native range result in lower fitness as estimated using population growth rates (Pantoja, Paine, & Vallejo-Marin, 2018). The effects of admixture may be particularly strong on invasive species with a widespread, highly diverse native distribution, such as *M. guttatus*. Native populations that occur over large, biogeographically diverse areas may serve as reservoirs of genetic and ecological variation. This wide range of ecogeographic variation may facilitate the colonisation of new regions in the introduced range and potentiate the effects of subsequent introductions and admixture on the performance and adaptive potential of invasive populations.

## Acknowledgements

We thank John Willis and current and former members of his lab, including David Lowry and Kevin Wright, for providing access to North American seed material collected over many years, and to the Botanical Society of Britain and Ireland for their continued support locating UK *Mimulus* populations. Arielle Cooley kindly provided seed material from Chilean populations of *M. luteus var. variegatus*. We are very grateful to Claudia Buser and John Bailey for providing the New Zealand material, and Nils Bunnefeld, Anna Maria Fosaa and Símun Arge for their help while collecting *Mimulus* in the Faroe Islands. We thank Oregon Genomics (University of Oregon) for sequencing services, the University of Stirling Controlled Environment Facility for access to plant growth facilities, and Sophie Webster for help in the laboratory. Computer time for the ABC analysis was provided by the computing facilities MCIA (Mésocentre de Calcul Intensif Aquitain) of the Université de Bordeaux and of the Université de Pau et des Pays de l’ Adour. LY was supported by the European Research Council (ERC) under the European Union’ s Horizon 2020 research and innovation programme [grant number ERC-StG 679056 HOTSPOT], via a grant to LY. This project was made possible by a grant from the Global Exploration Fund, Northern Europe from National Geographic (GEFNE164-15) to MVM, JRP and SMI-B, and a grant from the Natural Environment Research Council (NERC; NE/J012645/1) to MVM. We thank all the people who helped us during fieldwork in Alaska, particularly Suzi Golodoff (Unalaska Island) and Stacy Studebaker (Kodiak Island) for providing their exceptional knowledge of the local flora, and Roger Topp (U. Alaska, Fairbanks/Museum of the North) who documented the expedition with his outstanding photographs and video.

## Data Accessibility

Genotype data will be made available upon publication as VCF files in a public repository (DATAStorre, U. Stirling). Location data of sampled populations is available in the Supplementary Materials. Herbarium specimens of newly collected material in Alaska is deposited at the ALA herbarium.

## Author contributions

MVM, JRP, SMIB, JF and ADT designed the research. MVM, JRP, JF, SMIB, MCR and MvK collected material. MVM, ADT, and OL analysed the data. MVM, JF, LY, ADT and JRP wrote the manuscript with input from all the authors.

## Supplementary Material

**Table S1**. Populations sampled and sequenced. Taxon: gut = *M. guttatus*; gut4x = tetraploid *M. guttatus*, lut = *M. luteus*; rob = *M. x robertsii*; per = *M. peregrinus*, gla = *M. glabratus*. Region: ak = Alaska; nam = western North America; enam = eastern North America; fo = Faroe Islands; uk = United Kingdom; eur = continental Europe (Germany); sam = South America; nz = New Zealand. Life history: A = annual; P = perennial; NA = not available.

**Table S2**. Posterior estimation of the demographic parameter of model E4. the introduced effective population size over current UK effective population size N0, divided by the time of first introduction to UK t0a).

**Figure S1**. Map of North America showing five groups of native *M. guttatus*. Groups were estimated using the global data set by *kmeans* clustering (k=8). Red = South group; yellow = North group; dark yellow = Coastal group; Blue = North Pacific group; orange = Aleutian group.

**Figure S2**. Population genetic structure of native and introduced populations of *Mimulus guttatus* inferred in a Bayesian approach using *fastStructure* (K=2 to K=8). For this analysis, all populations were limited to a maximum of 3 individuals per population. Individuals within geographic regions are arranged by cluster membership. Alaska (native), Western North America (native); ENA = Eastern North America (introduced); GER = Germany (introduced); FO = Faroe Islands (introduced); NZ = New Zealand (introduced); United Kingdom (introduced).

**Figure S3**. Demographic reconstruction of the origin of invasive populations of *Mimulus guttatus* in the United Kingdom using Approximate Bayesian Computation (ABC). The scenario shown here (E4) was selected by hierarchical testing increasingly complex models starting with a single origin of extant UK populations. The model shown here, suggests a first introduction from the Aleutian Islands followed by additional introductions from other parts of the native range of *M. guttatus*.

